# Patterned substrates modulate growth and dynamics of 3D cellular systems

**DOI:** 10.1101/632034

**Authors:** Michael J. Fanous, Yanfen Li, Mikhail E. Kandel, Kristopher A. Kilian, Gabriel Popescu

## Abstract

The development of 3D cellular architectures during development and pathological processes involves intricate migratory patterns that are modulated by genetics and the surrounding microenvironment. The substrate composition of cell cultures has been demonstrated to influence growth, proliferation, and migration in 2D. Here we study the growth and dynamics of mouse embryonic fibroblast (MEF) cultures patterned in a tissue sheet which then exhibits 3D growth. Using gradient light interference microscopy (GLIM), a label-free quantitative phase imaging approach, we explored the influence of geometry on cell growth patterns and rotational dynamics. We apply, for the first time to our knowledge, dispersion-relation phase spectroscopy (DPS) in polar coordinates to generate the radial and rotational cell mass-transport. Our data show that cells cultured on engineered substrates undergo rotational transport in a radially independent manner and exhibit faster vertical growth than the control, unpatterned cells. The use of GLIM and polar DPS provides a novel quantitative approach to studying the effects of spatially patterned substrates on cell motility and growth.

## 1. Introduction

The mechanical properties of a cell’s microenvironment play a role in guiding cellular functions, including cell migration and growth patterns[1–3]. Cells sense the extracellular matrix (ECM) via cell surface receptors which then propagate the mechanochemical signals into the cell to influence cellular functions downstream[4]. Cellular behavior can thus be modulated by mechanical[5], biochemical[6, 7], or thermal[8] stimuli. Substrate engineering provides a reproducible approach to study these mechanochemical cues including the ability to control the boundary mechanics and geometry of cells and tissues[9–11]. Micropatterning of large populations of cells produces a gradient of force spatially organized within the pattern which then leads to differing patterns of cellular function[12], including cellular differentiation[13, 14] and migration[15]. Patterning the geometry of culture substrates causes mutations to a cell’s intrinsic makeup, which is phenomenon that remains insufficiently understood[10, 15–18]. The 3D dynamics of biological specimens, such as organoids, can be diverse, complex, and challenging to assess precisely[19, 20]. Though several studies have reported on the effects of cell cultures[21], no technique, to our knowledge, has been used to simultaneously tackle quantitative growth and mass transport.

Typical methods to characterize complex biological specimens involve exogenous contrast agents, such as fluorescence dyes[22, 23]. These artificial inserts affect the cell’s original composition and can lead to damage via phototoxicity[24]. A further limitation with fluorescent modalities is the photobleaching effect that restricts imaging duration[25].

Quantitative phase imaging (QPI)[26] is a *label-free* approach that has emerged as a powerful alternative to measuring biological phenomena quantitatively[27]. QPI’s key feature is that it can image cells nondestructively[28, 29]. The recurrent drawback with laser instruments, however, has been the artifact of speckles resulting from coherent light sources[30–32]. During the past decade, several white light based methods have been devised to obtain speckle-free images[33–37]. In particular, Spatial Light Interference Microscopy (SLIM)[26] and Gradient Light Interference Microscopy (GLIM)[37] have been used successfully to study the 3D morphology of various sample types, such single cells[38, 39], embryos[27], red blood cells[40], and neural networks[41]. Very recently, SLIM has been used for high-throughput single cell weight phenotyping in biomass producing cell populations[42]. GLIM is particularly well suited for subduing multiple scattering backgrounds and for effectively reducing out-of-focus light. Both systems employ a spatial light modulator to combine phase shifting and common path interferometry and generate quantitative phase information. SLIM and GLIM are implemented as add-ons to customary phase contrast[43] and differential interference contrast (DIC) microscope stands[44], respectively. Dispersion-relation phase spectroscopy (DPS)[45] has been developed as an analysis method for phase images to inspect diffusion and advection signatures in cellular systems[41, 46–48].

In this paper, we use GLIM in conjunction with a modified version of the DPS technique, to investigate the growth and rotational dynamics of MEF cultures on patterned substrates as they grow into 3D aggregates, approximating spheroidal architecture. We observe that substrate patterning induces the cells to exhibit greater vertical growth and that the cells undergo rotational fluctuations in a plane parallel to the substrate. Our methods will likely facilitate a deeper understanding of 3D migration and proliferation processes that underlie cellular assembly into tissue-mimetic structures, during morphogenesis[49].

## 2. Methods

Unless otherwise mentioned, all materials were acquired from Sigma-Aldrich. Tissue culture plastic ware was purchased from VWR. Glass coverslips were purchased from Fisher Scientific. Cell culture media reagents were purchased from Gibco.

### 2.1 Cell Culture

Mouse embryonic fibroblasts (MEFs) were obtained through a generous donation from Dr. Quanxi Li of the Department of Comparative Biosciences, University of Illinois at Urbana-Champaign. The cells were cultured in glucose (5g/mL) DMEM supplemented with 15% fetal bovine serum (FBS, Invitrogen) and 1% penicillin/streptomycin. Cells were passaged at 80% confluency with 0.5% Trypsin: EDTA and the growth medium was replaced every 3 days. For imaging, cells were seeded at ~200,000 cells/cm in a 6 well glass bottom plate (P06-20-1.5-N) and were imaged over a duration of 100 hours every 30 minutes with an acquisition rate of 6 frames/s. The microscope housed an incubator unit to sustain the cells at 37 degree Celsius, and 5% CO_2_. Wells either had patterned or normal substrate configurations, and each field of view was imaged by a 10-layer z-stack of 10 μm increments. Surface plots and videos were generated through maximum projection renderings. The heights of the sample were determined by computing the peak maximum values of normalized phase gradients in each frame of the z-stack.

### 2.2 Gel Preparation

10 kPa polyacrylamide hydrogels were produced as previously described to mimic the stiffness of an MEF’s natural bioenvironment[50]. In brief terms, a mixture of 5% polyacrylamide and 0.15% bis-acylamide was fabricated and then combined with 0.1% Ammonium Persulfate (APS) and 0.1% Tetramethylenediamine (TEMED)[51]. Solutions were placed onto a hydrophobically prepared glass slide (Rain-X) and sandwiched between an aminopropyltriethoxisilane (APTES)- sinalized glass coverslip. Gels were removed from the coverslip after polymerization and immersed in 55% hydrazine hydrate (Fisher) for one hour, and then washed with 5% glacial acetic acid for one hour.

### 2.3 Gel Pattering

Circular SU-8 silicon masters were fashioned using photolithography. To produce stamps, Polydimethysiloxane (PDMS, Polysciences, Inc) was polymerized over the silicon masters[11]. 25 μg/ml fibronectin was incubated with Sodium Periodate for 45 minutes and incubated on top of PDMS stamps for 30 minutes. The stamps were then air dried and applied to the hydrogels to create the desired patterns.

### 2.4 Gradient Light Interference Microscopy

Measurements were performed using GLIM, consisting of an inverted differential interference contrast (DIC) microscope (Axio Observer Z1, Zeiss, in this case) and an add-on module (*CellVista GLIM Pro*, Phi Optics, Inc). GLIM generates quantitative phase gradient images of the sample with high depth sectioning capability. Quantitative phase methods often use coherent light sources that hamper the contrast of the images due to scattering artifacts[37]. GLIM resolved this problem by using low-coherence interferometry with white light, enabling extremely sensitive measurements. GLIM is also label-free, which allows imaging for long durations without imposing deleterious conditions on the cells.

## 3. Polar DPS

To study the translational and radial components of cellular mass transport, we used the dispersion-relation phase spectroscopy (DPS) technique[45] in polar coordinates. This method is effective at distilling spatiotemporal dynamic information from a time-lapse sequence of phase images. It does not require any manual tracing or labelling and enables automatic computations[41]. In DPS, the dispersion relation of the motile system is evaluated, combining the spatial and temporal frequencies. The nature of the dispersion curve informs on the type of transport, whether it is diffusive or deterministic. For our analysis, we first convert the GLIM images into polar coordinates and then perform DPS as previously described[41]. However, instead of taking an azimuthal average of the power spectrum, we evaluated both the rotational and radial dynamics separately. The dry mass-density fluctuations can be described via an advection-diffusion equation as follows,

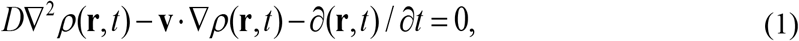

where **r** = (*r,θ*) indicates the 2D coordinate, **v** is the advection velocity, and *D* is the diffusion coefficient. Using this equation, we obtain the temporal autocorrelation function computed at every spatial frequency, **r** = (*r,θ*) and temporal delay, *τ*, defined as 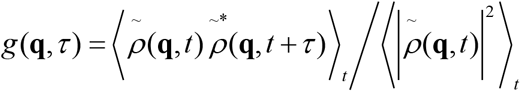. Here, 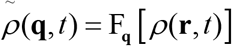 is the 2D spatial Fourier transform of the dry mass density. It has been shown previously that one can calculate *g*(**q**, *τ*) directly from the phase gradient ▽*ϕ* instead of the phase itself *ϕ*[37], which yields

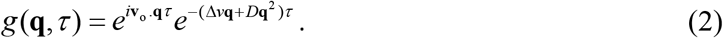

where **v**_o_ is the mean and Δ*v* the standard deviation of the velocity distribution. At each rotational or translational spatial frequency, one can fit the measurement of *g*(**q**, *τ*) to compute Δ*v_θ_* and Δ*v_r_*. The decay rate of *g*(**q**, *τ*) is governed by

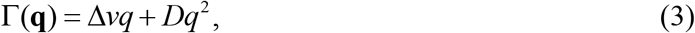

where Δ*v* corresponds to the active transport and *D* refers to diffusion.

## 4. Results

### 4.1 Effects of circular substrate patterning on 3D cell growth

To explore how geometric constraints influence the spatiotemporal development of large populations of cells, mouse embryonic fibroblasts (MEFs) were cultured on both patterned and control (non-patterned) polyacrylamide hydrogel matrices of 10kPa stiffness (Fig. 1). Cells cultured on fibronectin coated polyacrylamide can move readily and proliferate without impediment[51]. We chose to focus on MEFs because these cells are poised to proliferate and undergo morphogenetic transformations during normal development of the embryo. Polyacrylamide hydrogels were chosen because they can be fabricated to encompass numerous normal and pathological mechanical properties[52, 53]. Previous research relied on qualitative or fluorescent methods that do not quantify cellular mass accurately[54].

**Figure 1.**
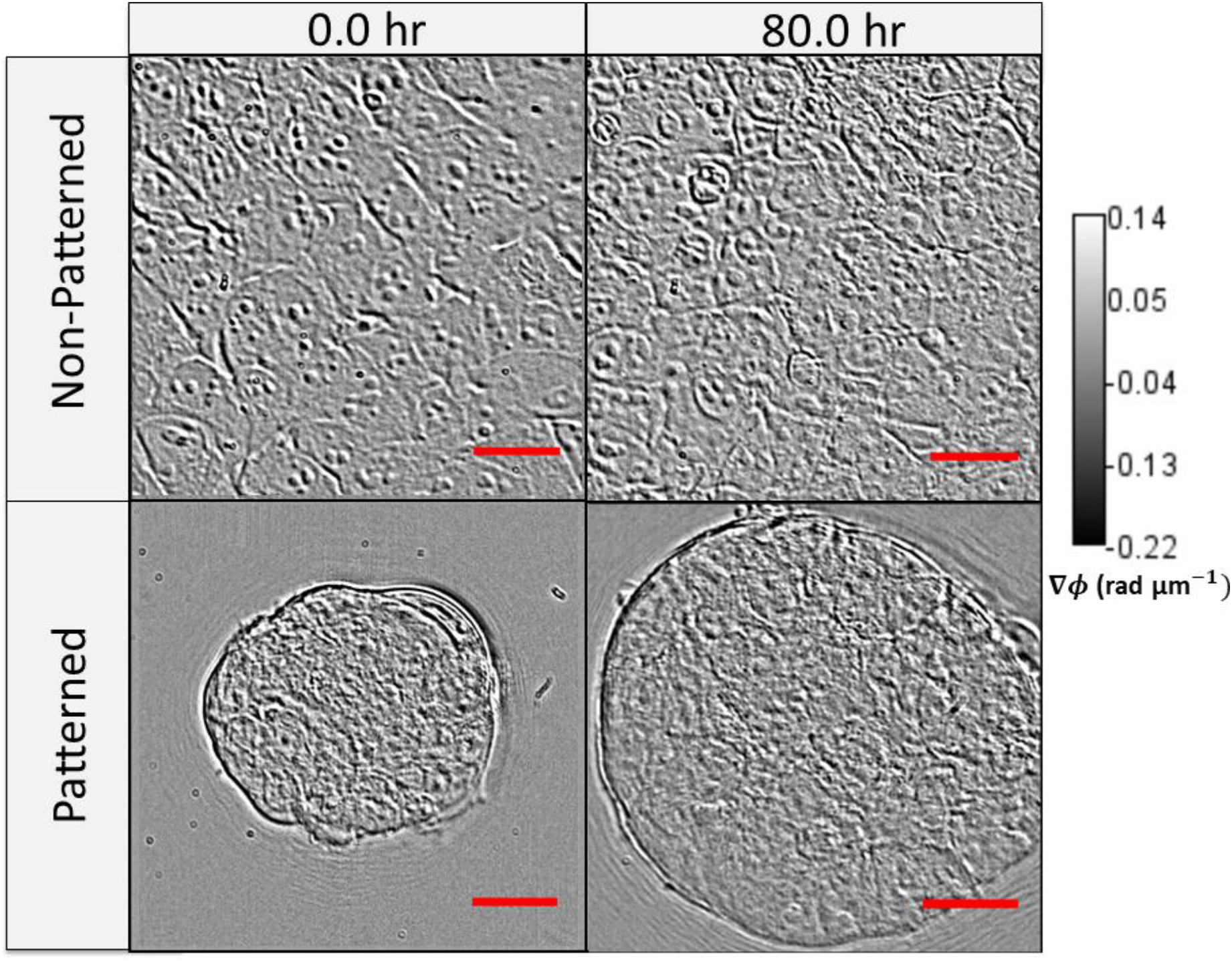
Gradient light interference microscopy (GLIM) images of mouse embryonic fibroblasts cultured on non-patterned and patterned substrates (polyacrylamide gels - 10 kPa), shown before and after 80 hours of growth. Scale bar: 30 μm.

In order to quantify growth and transport characteristics, we seeded MEFs on patterned and control hydrogel substrates and imaged them with our GLIM system. The samples were imaged every thirty minutes for 100 hours. GLIM performs label-free measurements of optically thick samples and outputs quantitative phase gradient maps. We found that the patterned cells exhibit significant expansion and elevation as compared to the cells cultured on standard substrates (Fig. 2). By computing the cell height through a maximum average computation, we found that patterned cells reached a height of 20 μm above their unpatterned counterparts. This behavior is captured quantitatively in Fig. 3. These data are consistent with previous work showing that designed substrate geometries induce differential and more robust growth tendencies[11].

**Figure 2.**
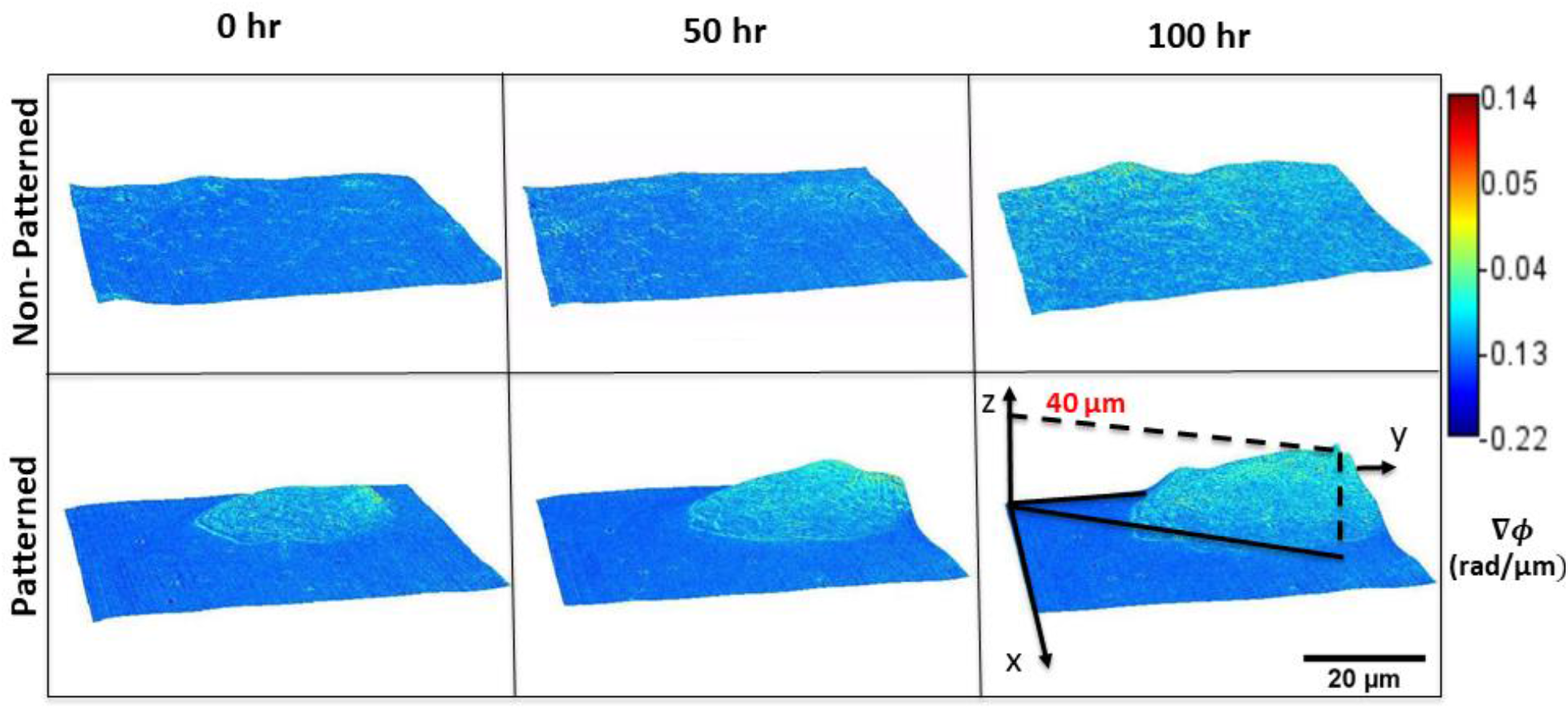
Gradient light interference microscopy (GLIM) images of mouse embryonic fibroblasts cultured on non-patterned and patterned substrates (polyacrylamide gels - 10 kPa), shown at 50 hour intervals. The cellular cluster on patterned substrates exhibits lateral growth while rising and rotating about the vertical axis.

**Figure 3.**
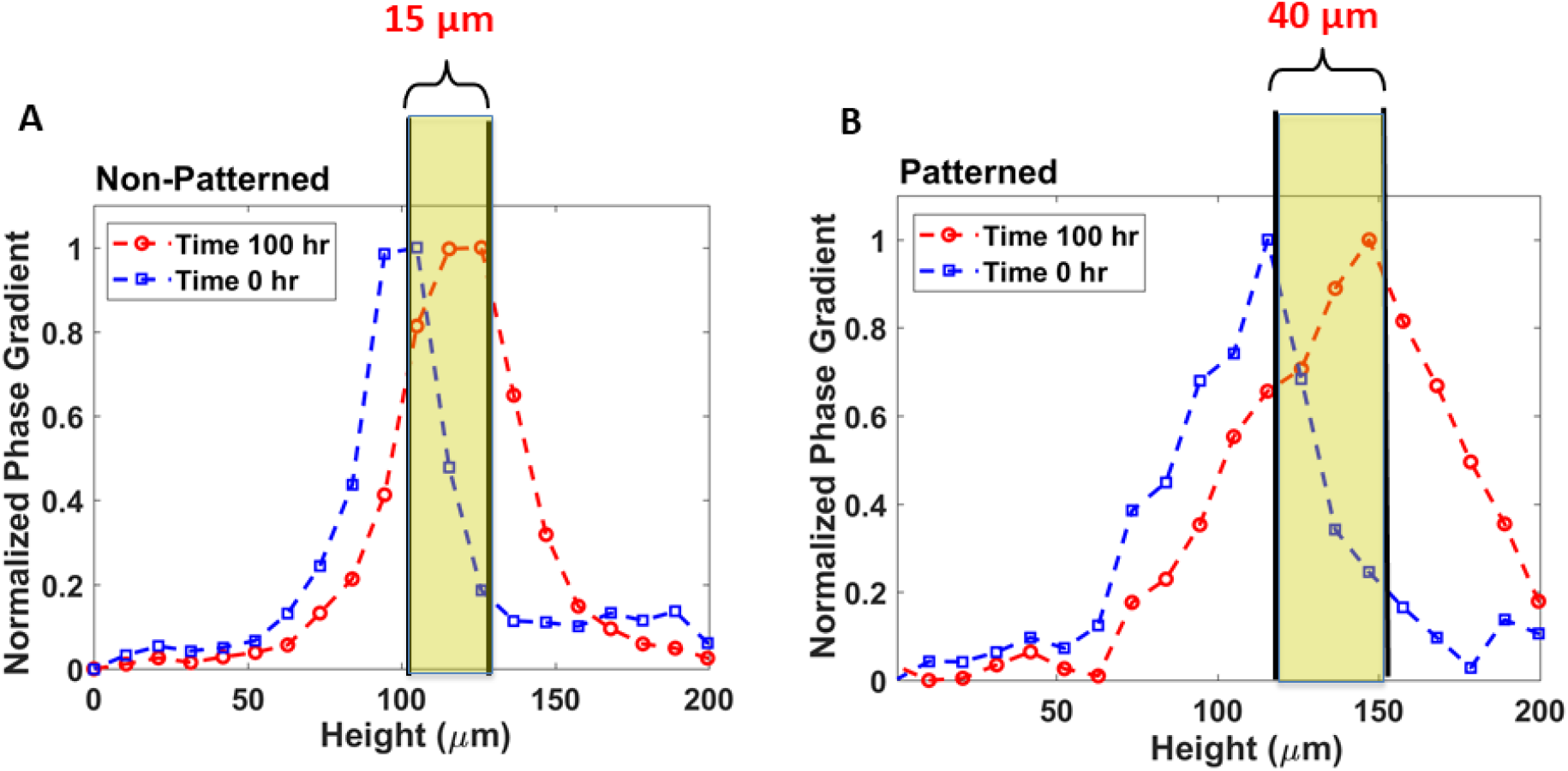
Total substrate height determined from peak maximum values of normalized phase gradients for non-patterned substrates indicate a 15 μm height increase over 100 hours (A). Total substrate height determined from the peak maximum values of normalized phase gradients for patterned substrates indicate a 40 μm height increase over 100 hours (B).

### 4.2 Patterned cells display inhomogeneous radial and rotational mass transport

We applied a novel technique for investigating the dynamic properties of cellular transport. We calculated the standard deviation of the *angular* and *radial velocity* distribution for MEFs on patterned and unpatterned hydrogels. These parameters were obtained from the slope of the decay rate within the angular frequency range of (0, 10) rad/rad, for rotational motion, and (0, 0.8) rad/μm for translational motion (Fig. 4). These intervals correspond to structures as large as the field of view, down to measurements as small as 0.314 rad and 3.9 μm, respectively. Figure 5 shows an example of plots of polar DPS curves for both rotational and translational action. Curves of the rotational measurement show slopes in the range of 0.4-0.6 rad/hr and curves of the radial DPS measurement have slopes in the range of 10-12 μm/hr, in the frequency range (q_*θ*_ < 10 rad/rad, q_*R*_ < 1rad/ μm). These findings are compatible with previous work indicating coordinated rotation of large population of cells when patterned in circular shapes[15] and with migration results of melanoma cells in our previous work[11].

**Figure 4.**
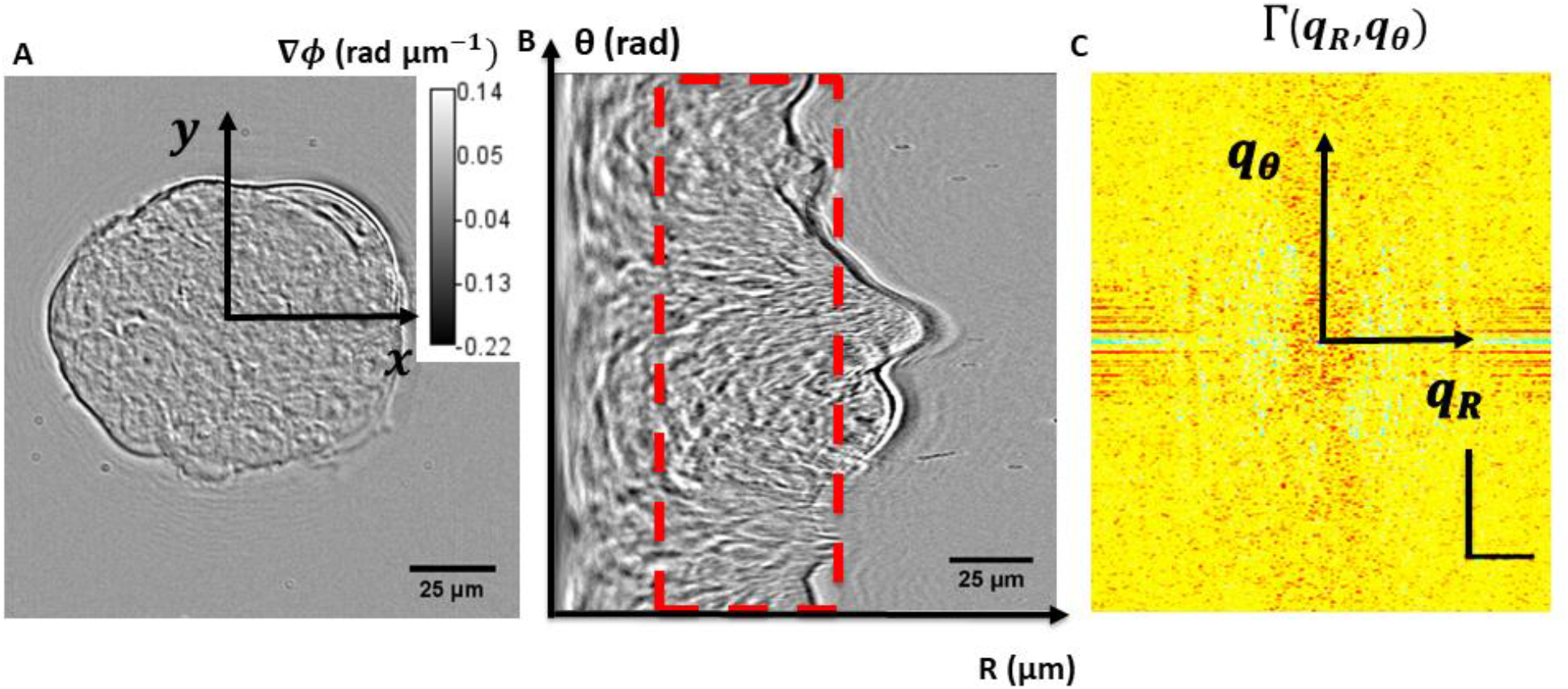
GLIM image of mouse embryonic fibroblasts cultured on a patterned substrate (A), and the corresponding polar transformation of the image, with an angular vertical dimension and a horizontal radial dimension, and showing the region or “ring” (red dashed rectangle) that was used in the computation (B). Decay rate vs spatial mode (angular and radial) associated with polar GLIM images (horizontal scale bar: 20 rad/μm; vertical scale bar: 20 rad/rad) (C).

**Figure 5.**
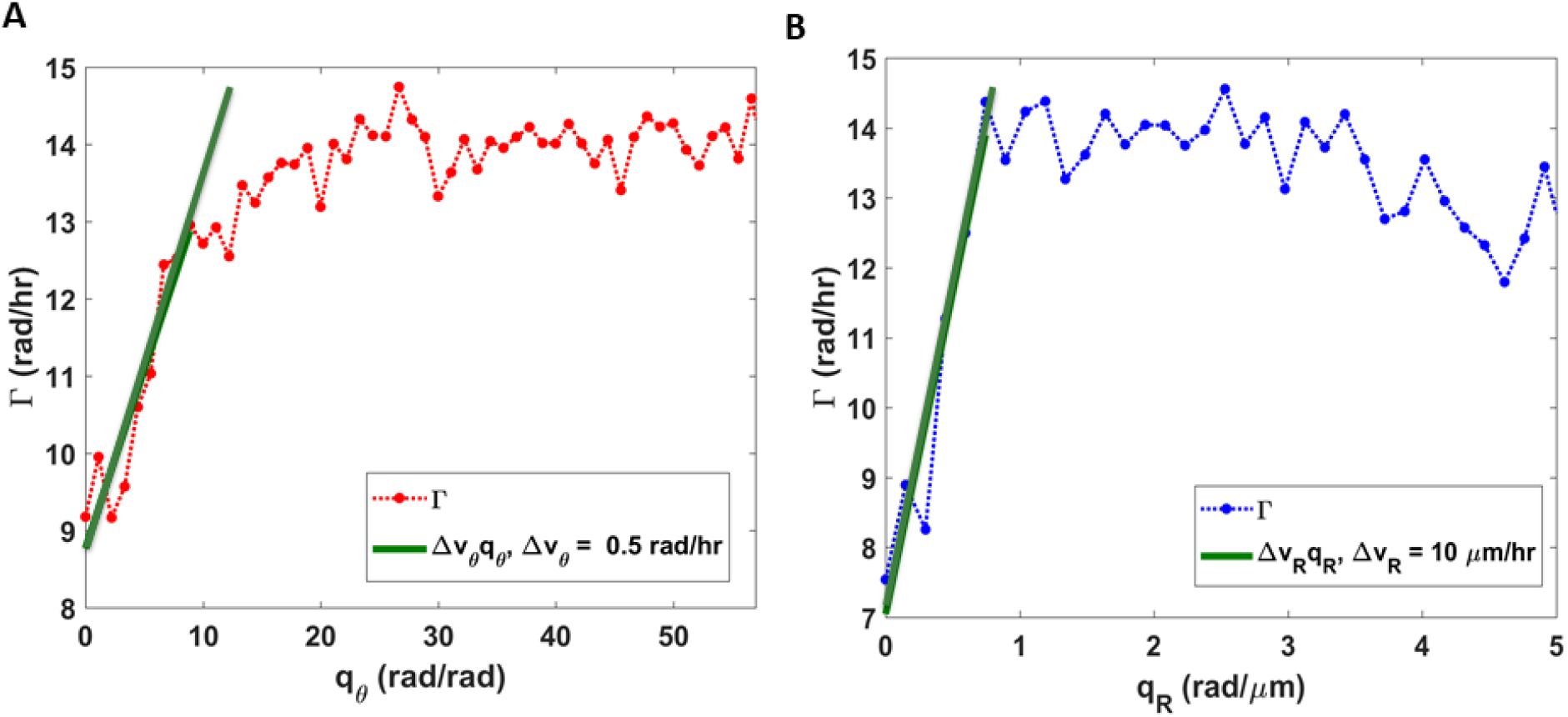
Angular decay rate with a slope of the linear fit down to 0.34 radians, indicating the angular velocity distribution width about a certain diameter (56 μm) (A). Radial decay rate with a slope of the linear fit down to 3 μm, indicating the radial velocity distribution width around the centroid of the patterned substrate (B).

### 4.3 Testing the radial homogeneity and angular isotropy of cellular mass transport

Next, we asked the question whether the angular velocity distribution changes with radial position. Such variations would indicate that the cells produce shear stress between different concentric layers. Figure 6A shows that this is, in fact, not the case. The rotational velocity distribution width exhibits small fluctuations with the radius Δ*v_θ_* = Δ*v*_*θ*_0__ ± *σ_θ_* = 0.16 ± 0.06 *rad* / *hr*. The translational velocity distribution with respect to angle informs about the anisotropy of mass transport. Figure 6B indicates that there is no significant monotonic variation with angle: Δ*v_r_* = Δ*v*_*r*_0__ ± *σ_r_* = 2 ± 1.5 *μm* / *hr*. These results hint at collective and harmonious cellular behavior that may imply emergent properties engendered through substrate patterning.

**Figure 6.**
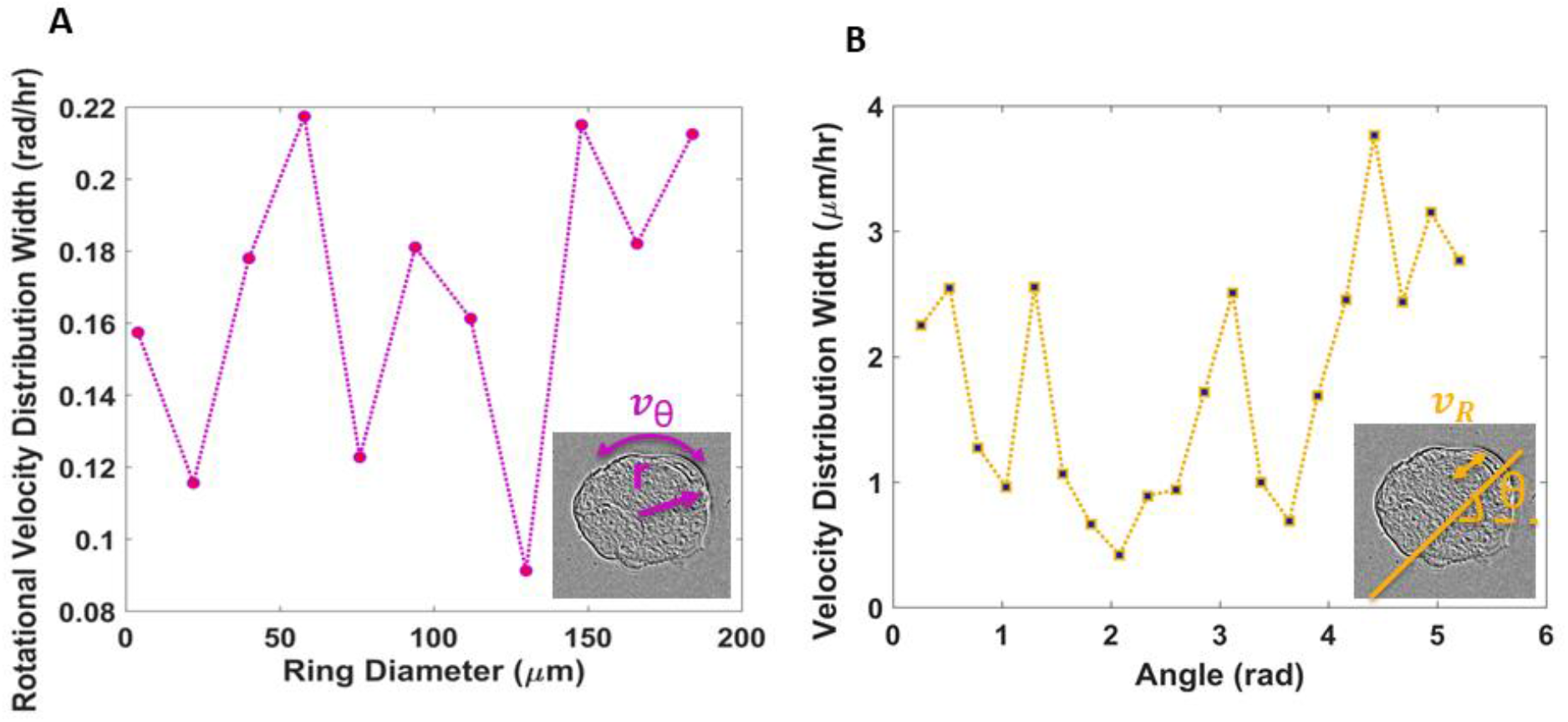
Rotational velocity distribution width of rings of the patterned substrate at varying inner diameters, with a ring thickness of 28 μm (A). Velocity (radial) distribution width at different angles (B).

## 5. Conclusion

In this paper, we used label-free, high-throughput imaging to unravel unique growth and migration trends in 3D cellular systems, which are directed by initial conditions of the substrate mechanics and geometry. The combination of quantitative imaging with DPS in polar coordinates was used to quantify growth disparities caused by matrix confinements and to reveal specific cell movement patterns. MEFs cultured on engineered substrates exhibited greater out-of-plane growth. Cells situated along the edge of the culture disc showed angular velocity distributions similar to those closer to the center, which suggest that there is no significant shear stress along the radius. Rotational transport appears to be isotropic, consisted with the symmetry of the pattern.

We have demonstrated how combining quantitative phase imaging with specially designed substrates can determine variations in cell dynamics throughout 3D cultures, which may prove beneficial in studying spheroids, organoids, and microtumors. Future research will involve employing this methodology to study asymmetric substrate geometries and correlation between the dynamic parameters and cellular metastatic potential.

## Acknowledgements

This work was supported by the National Institute of General Medical Sciences (NIGMS) grant GM129709, the National Science Foundation (NSF) grant CBET-0939511 STC, DBI 14-50962 EAGER, IIP-1353368, as well as the (NSF)-funded Miniature Brain Machinery (MBM) program.

